# Effects of electroacupuncture on the expression of hypothalamic neuropeptide Y and ghrelin in pubertal rats with polycystic ovary syndrome

**DOI:** 10.1101/2021.10.25.465707

**Authors:** Yang Li, Wang Zhi, Dong Haoxu, Wang Qing, Cheng Ling, Yi Ping, Huang Dongmei

**Affiliations:** Institute of Integrated Traditional Chinese and Western Medicine, Tongji hospital, Huazhong University of Science and Technology, Wuhan, China; Department of Integrated Traditional Chinese and Western Medicine, Tongji hospital, Huazhong University of Science and Technology, Wuhan, China; Department of Rehabilitation Center of Wuhan Puren Hospital, Affiliated Hospital of Wuhan University of Science and Technology, Wuhan, China; Department of acupuncture and moxibustion, east hospital, tongji university, Shanghai, China

**Keywords:** Polycystic ovary syndrome, adolescence, electroacupuncture, hypothalamus, neuropeptide Y, ghrelin

## Abstract

**Background:** Polycystic ovary syndrome often starts in puberty, and its pathogenesis is not clear. This study aimed to explore the pathogenesis of pubertal polycystic ovary syndrome (PCOS) and assess the therapeutic effect of electroacupuncture on pubertal PCOS.

**Methods:** Dihydrotestosterone (DHT) was used to induce rat models of pubertal PCOS. pubertal rats with PCOS were randomly divided into a model group (M), an electroacupuncture group (EA), and a sham acupuncture group (SA). Age-matched normal rats were regarded as normal controls (N). Rats were treated with EA or SA five times a week for 25 minutes during their 6^th^–7^th^ week. At the end of the experiment, we observed any changes in ovarian morphology; detected levels of metabolic indices in serum, the hypothalamus and pancreas.

**Results:** EA significantly improved estrous cycle disorders and the ovarian polycystic morphology in pubertal rats with PCOS, but SA only improved disorders of the estrous cycle. The serum levels of insulin, neuropeptide Y(NPY) and fasting blood glucose(FBG) increased significantly, while the serum levels of ghrelin(GHRL) decreased in the model group. After treatment with EA, the levels of NPY and FBG went into decrease, whereas the levels of GHRL and insulin increased. There was few differences in the hypothalamic expression of galanin (GAL), galanin-like peptide (GALP) and ghrelin receptor(GHSR) between the four groups. The upregulation of NPY mRNA and neuropeptide Y2 receptor(NPY2R) mRNA and the downregulation of GHRL protein and mRNA in the hypothalamus, and the increased expression of NPY and NPY2R as well as the decreased expression of GHRL in the arcuate nucleus (ARC) can be rescued by EA. But, surprisingly, SA seem to make no difference to the levels of FBG and insulin, and the protein expression of ghrelin in the hypothalamus and ARC. Co-expression of kisspeptin and GHSR, and co-expression of gonadotrophin releasing hormone(GnRH) and NPY2R were observed in ARC. No differences were found between groups in protein of GAL, GALP and GHRL expression in the pancreas. Neither EA nor SA can attenuate the upregulated kisspeptin protein expression in the pancreas of PCOS model rats.

**Conclusions:** EA and SA improved the symptoms of pubertal PCOS rats, and the mechanism might be associated with regulating hypothalamic NPY and ghrelin levels.

## Introduction

Polycystic ovary syndrome (PCOS) is a common reproductive disease, leading to endocrine disorders, metabolic disorders, and infertility in women. Patients with PCOS usually seek medical attention during their childbearing years due to infertility. However, most patients also show obvious clinical symptoms in adolescence [1], such as hyperandrogenism, oligomenorrhea, amenorrhea, and metabolic disorders. The elevated luteinizing hormone(LH) pulse frequency and increased activity of the gonadotropin-releasing hormone (GnRH) neural network are typical characteristics among women with PCOS[2–4]. Meanwhile, GnRH neuronal network generates pulse and surge modes of gonadotropin secretion critical for puberty[5]. Girls treated with GnRH analog drugs (GNRHa) have a significantly increased risk of PCOS during adolescence[6]. Pubertal rats with PCOS showed neuroendocrine dysfunction, characterized by hypersecretion of LH and kisspeptin and elevated release of GnRH from hypothalamus[7]. This elevated hypothalamic GnRH output is likely to stem in part from changed of the metabolic state of the body. Neuropeptide Y(NPY), ghrelin(GHRL), galanin (GAL) and galanin-like peptide (GALP) have been proposed to be the candidates conveying metabolic status to the GnRH neuronal network in animals.

NPY, known as an orexigenic neuropeptide, is strongly implicated in linking metabolic cues with fertility regulation[8]. NPY were observed to be increased continuously from birth to the time of vaginal opening, and a surge in portal levels of NPY which preceded the prepubertal LH surge[9]. After NPY treatment of rhesus monkey hypothalamic culture, GnRH increased 20-30 minutes later[10]. Ghrelin, an orexigenic hormone, is mainly produced by stomach and released to circulation. However, ghrelin was be found to be also expressed in hypothalamus and inhibit GnRH secretion[11]. GAL, synthesized by a subgroup of GnRH neurons in hypothalamus[12], is a neuropeptide widely distributed in the central and peripheral nervous systems. Research has shown that GAL could directly stimulate LH secretion at the pituitary level and enhance GnRH stimulated LH secretion[13, 14]. GALP is a recently identified hypothalamic peptide, localized in the arcuate nucleus(ARC), which seems to stimulate hypothalamus and GT1-7 cells (a GnRH neuron cell line) to release GnRH[15]. Levels of circulating GAL during puberty have been discovered to be significantly higher than those in pre-puberty in patients with type 1 diabetes mellitus, while levels of ghrelin were significantly lower[16]. Central GALP treatment has been able to rescue the onset of puberty in food-restricted weanling rats[17].These suggest involvement of NPY, GHRL, GAL and GALP in reproductive development and the onset of puberty.

Abnormal metabolic disorders, such as insulin resistance, glucose intolerance, hyperlipidemia, and obesity, are found to be closely related to PCOS and appear during puberty. In several studies, abnormal expressions of metabolic molecules (for example, the higher level of NPY and GAL, whereas the lower ghrelin levels) have been found in patients with PCOS[18–20]. However, how those metabolic molecules change in adolescent PCOS is still unknown.

Electroacupuncture (EA) has a benign regulating effect on neuroendocrine levels in organs. Repeated low-frequency EA inhibited the expression of GnRH and NPY in the hypothalamus of early adolescent Sprague Dawley (SD) female rats[21]. It has also been reported that EA can inhibit the secretion of GnRH and the expression of androgen receptors in the hypothalamus of PCOS rats[22]. However, there is little research on the effect of EA on PCOS in puberty. The aim of this study was to investigate the alteration of metabolic molecules in adolescent PCOS and whether EA plays a role in adolescent PCOS through the modulation of these metabolic molecules.

## Materials and methods

### Animals

Female SD rats weighting 250 g and male SD rats weighing 300 g were purchased from Beijing Weitonglihua Experimental Animal Technology Co., Ltd, Beijing, China (Animal certificate: SCXK no. 2016-0006). All rats were fed ad libitum in experimental animal center of Huazhong University of Science and Technology, Wuhan, China (Demonstration of Laboratory Animal Facilities in Hubei Province, SYXK 2016-0057). Animal care and experimental procedures comply with the ARRIVE guidelines and were performed in accordance with the U.K. Animals (Scientific Procedures) Act, 1986 and associated guidelines, EU Directive 2010/63/EU for animal experiments (IACUC no. 2298).

Female rats were mated with males. From the 16^th^ day to the 19^th^ day of gestation, the rats were injected once a day subcutaneously with 0.3 ml sesame oil (Lot no. 1612426-2X12MG), containing 3 mg 5α-dihydrotestosterone (DHT, lot no. SZBD263XV). Method of establishing the puberal PCOS rat model is shown in Figure 1.

**Fig. 1.**
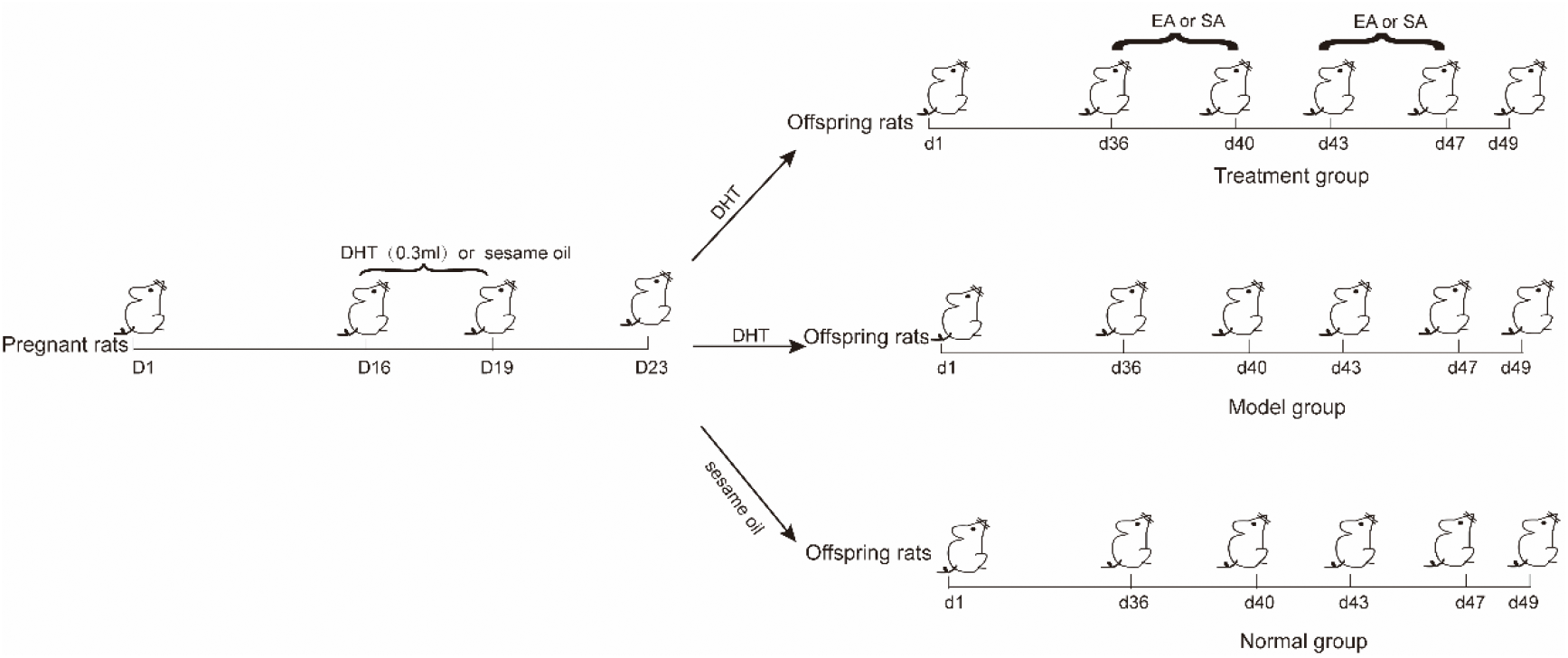
Process of establishing the puberal PCOS rats model.

### Vaginal smears and estrous cycle determination

The estrous cycles of the female rats were determined by saline vaginal smears. The female offspring of androgenized pregnant rats were smeared at 8:00 am from the 29^th^ day to the 49^th^ day. The stage of the estrous cycle was defined by the predominant cell type[23]. Estrous cycles are mainly divided into four stages including the proestrus, estrus, metestrus, and diestrus. Each stage is marked by the presence of different types of cells: proestrus by nucleated epithelial cells; estrus by keratinized acetous epithelial cells; metestrus by nucleated epithelial cells, keratinized acetous epithelial cells, and leukocytes; and diestrus by a large number of leukocytes. The PCOS model rats were characterized by disorders of the estrous cycle and prolonged diestrus.

### Experimental design and electroacupuncture administration

The modeled PCOS rats were randomly divided into four groups: the PCOS model group (M), the electroacupuncture group (EA), the sham acupuncture group (SA), and the normal group (N). The female offspring of normal female rats that were not injected with DHT were placed in the normal group.

In the EA group, the bilateral acupoints *sanyinjiao* (SP6) and *zusanli* (ST36), and the acupoint *qihai* (CV6) were chosen. Disposable, sterilized, stainless steel needles (Size: 0.16 × 25 mm; lot no. 491226; Wuxi Jiajian Medical Instrument, Wuxi, China) were inserted into those acupoints. The needles were connected to an electrical stimulator (Schwa-Medico GmbH, Ehringshausen, Germany), and the rats were then stimulated with EA at a low frequency of 2 Hz with a 0.3 ms pulse length and a 2-mA current, leading to gentle muscle quivering. The stimulation was repeated for 25 minutes once a day, five times a week, for two weeks from the 36^th^ to the 49^th^ day after birth.

In the SA group, the needling points were four non-acupoints. Two points were bilaterally three inches away from the *guanyuan* point (CV4). The other two points were bilaterally in the middle between the *zusanli* (ST36) and *yanglingquan* (GB34) points. Disposable, sterilized, stainless steel needles (size: 0.16 × 25 mm; lot no. 491226; Wuxi Jiajian Medical Instrument, Wuxi, China) were inserted subcutaneously to a depth of <5 mm and connected to an electrical stimulator (Schwa-Medico GmbH, Ehringshausen, Germany) but without increasing the intensity. The intervention period was the same as for the EA group.

The rats in the normal control group and the model group were fixed with special cloth bags for 25 minutes every day without any needling. Interventions for these groups took place five times a week for two weeks from the 36^th^ to the 49^th^ day after birth.

### Sample collecting

All the rats were decapitated on the 49^th^ day. First, blood samples were collected via the abdominal aorta after deep anesthesia. The rats were confirmed as dead after blood samples had been taken and before other operations could be continued. Brain, ovary and pancreas tissues were collected for analysis by western blot and RT-PCR testing. Brain tissues from a sample of rats were perfused with paraformaldehyde, and frozen sections were used in immunohistochemistry and immunofluorescence detection procedures.

### Ovarian morphology

The ovaries were fixed in 4% paraformaldehyde, embedded in paraffin and then cut longitudinally and consecutively into 4 μm slices. After every tenth slice, the sections were stained with hematoxylin and eosin and placed under an optical microscope so that their morphology could be observed.

### Determination of the serum levels of metabolic indicators

After fasting for 12 hours, the rats’ fasting blood glucose (FBG) was measured with a glucose meter (Roche, Basel, Switzerland). The serum levels of insulin (80-INSRT-E01), GHRL(EIAR-GHR-1) and NPY(EIAR-NPY-1) were determined using enzyme-linked immunosorbent assay kits.

### Western blotting

Hypothalamic tissues were dissolved in a RIPA solution containing protease and phosphatase inhibitors. Extracted protein samples were loaded onto an 8%–15% SDS polyacrylamide gel for electrophoresis and transferred onto PVDF membranes. After blocking in 5% non-fat milk at room temperature for 1 h, the membranes were incubated by primary antibodies (kisspeptin, catalog no. ab19028; GAL, catalog no. A1991; GALP, catalog no. A17445; GHRL, catalog no. A12581) at 4°C overnight. The secondary antibody was IRDye 800 cw Goat anti-Rabbit IgG (Lot no. 926-32111). The membranes were visualized with an Odyssey imager (LI-COR, USA).

### Quantitative real-time PCR (RT-qPCR)

Total mRNA was extracted from the hypothalamus using a TRIzol reagent according to the manufacturer’s instructions (Takara Biotechnology Co. Ltd, Dalian, China). RNA was reverse transcribed into cDNA using a cDNA reverse transcription kit (Catalog no. D006-1). The RT reaction was performed with mixed liquids containing cDNA, primer, and SYBR Green Master Mix on a real-time PCR instrument (Catalog no. D007-1). Data were normalized to β-actin mRNA and relative gene expression was calculated using the 2^−ΔΔCt^ method. Primer information is as follows (Table 1):

**Table 1:**
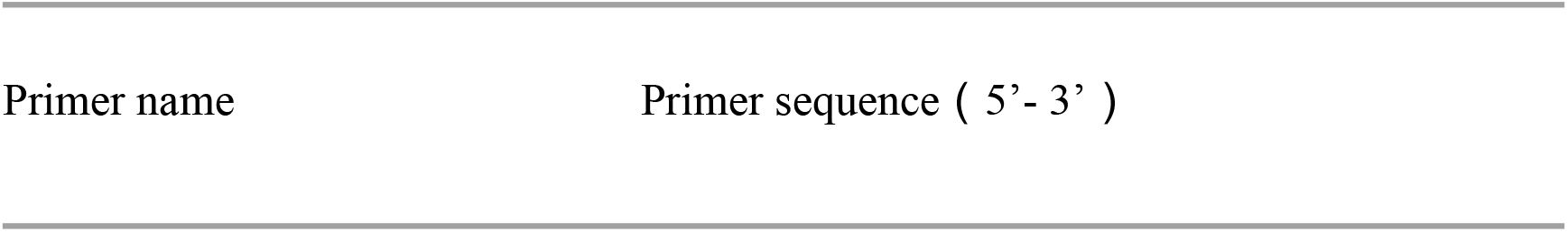
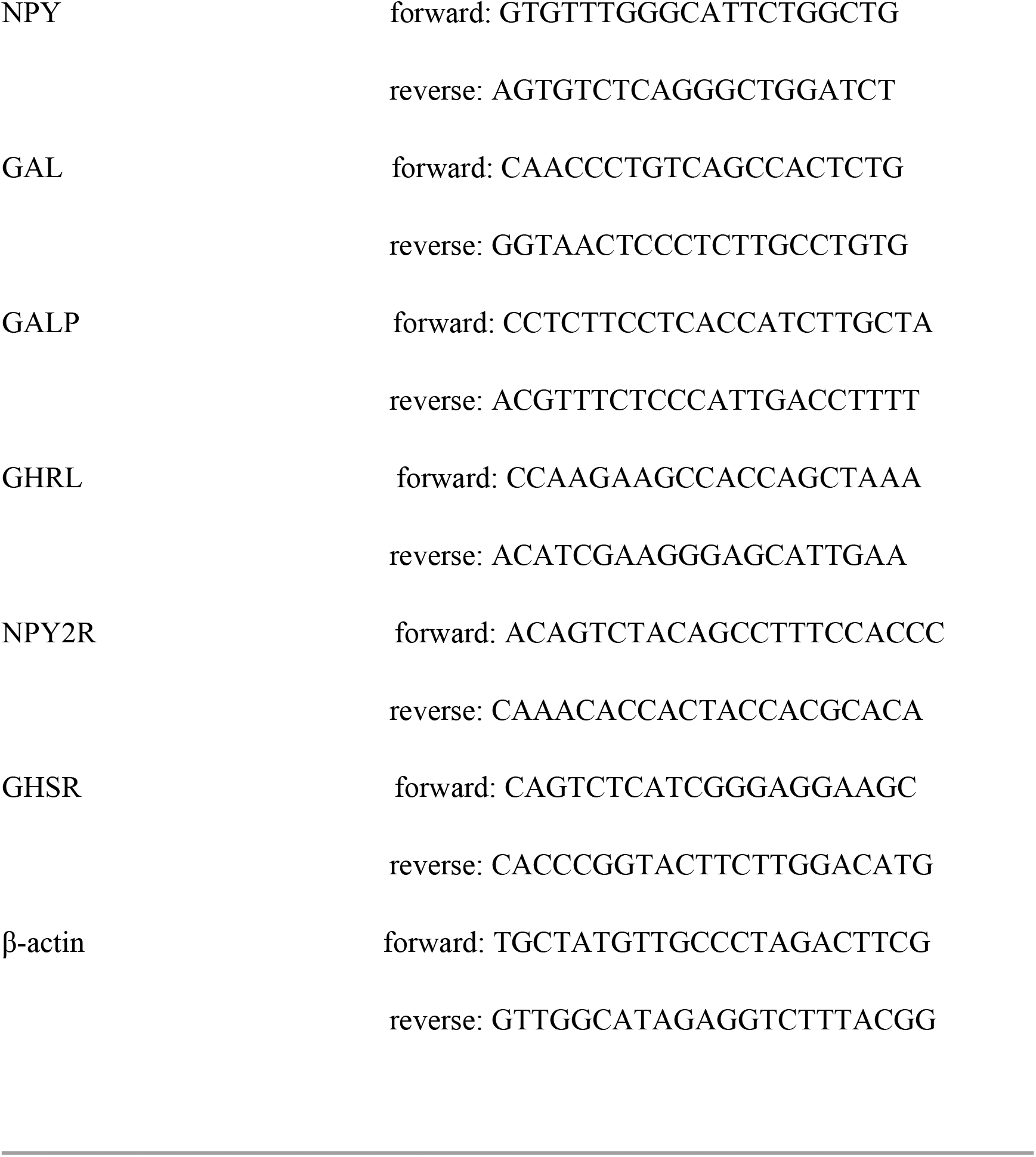
Primer Information

### Immunohistochemistry

The hypothalamus quickly separated from the brain for frozen section (8 μm) and the sections were placed in a refrigerator at −80°C. The frozen sections were covered successively with 4% paraformaldehyde for 10 minutes, hydrogen peroxide for 10 minutes, goat serum for 60 minutes, and finally with primary antibodies overnight at 4°C (NPY, catalog no. A3178; GHRL, catalog no. A12581; GHSR, catalog no. A1840; LEPR, catalog no. 20966-1-AP). After incubation with secondary antibodies ( catalog no. G1215) for 1 h at 37°C, the sections were stained with DAB reagent ( catalog no. G1212-200).

### Immunofluorescence

The frozen hypothalamus sections (8 μm) were kept at room temperature for 20 minutes, incubated with membrane-breaking fluid for 10 minutes, blocked with 10% donkey serum ( Catalog no. G1217)for 45 minutes, and then incubated with primary antibodies overnight at 4°C. The sections were washed with PBS three times (five minutes each time) before proceeding to the next step. Rabbit anti-NPY2R (Catalog no. DF5000) was mixed with mouse anti-GnRH (Catalog no. MAB5456), mouse anti-kisspeptin (Catalog no. MABC60) mixed with rabbit anti-GHSR (Catalog no. A1840), in order to test for co-localization. The marker used for cell nuclei was DAPI. The secondary antibodies were Alexa Fluor 594 Donkey anti-Rabbit IgG (ANT030) and Alexa Fluor 488 Donkey anti-Mouse IgG (ANT023).

## Statistical analysis

SPSS software (SPSS, version 21.0; Chicago, Illinois, USA) was used to analyze the data. The Kolmogorov–Smirnov test was applied to detect whether the data were normally distributed. The differences between the four groups were evaluated using the one-way ANOVA test for normally distributed data. The Kruskal–Wallis test was used to evaluate the differences in the skewed data. A value of P < 0.05 was considered to be statistically significant. Data are expressed as mean ± SEM.

## Results

### Estrus cycle and ovarian morphology

Disturbed estrous cycles could be observed in the model (M) group. When compared with the normal (N) group, the percentage of rats at estrus and proestrus was lower and the percentage of rats at diestrus was higher. After treatment with EA or SA, the percentage of rats at estrus significantly increased (Table 2). Although there was no statistical difference in the percentage of rats at proestrus and diestrus between groups M and EA, there was an opposite trend between the EA group and model group.

**Table 2.** The percentage of offspring at each stage of the estrus cycle during 4^th^ week to 7^th^ week in four groups

| Group | N | M | EA | SA |
| --- | --- | --- | --- | --- |
| Proestrus(%) | 22.02 ±2.69 | 11.31 ±1.79<sup>△</sup> | 13.10 ±1.19 | 11.90 ±1.80 |
| Estrus(%) | 19.05 ±1.27 | 13.69 ±1.40 | 17.86±1.49* | 18.45±1.40* |
| Mestestrus(%) | 19.05 ±2.54 | 15.48±1.96<sup>△</sup> | 13.69±1.40 | 18.45±2.45 |
| Diestrus(%) | 39.88 ±5.00 | 59.52±3.92<sup>△</sup> | 54.76±2.84 | 51.19±3.45 |

<sup>△</sup> stands for P < 0.05 versus Normal group; \* stands for P < 0.05 versus Model group;
Data are presented as means ± SEM;

HE-stained ovarian sections were observed under an optical microscope and the histological structure of the follicles was analyzed. As shown in Figure 2, Normal follicles at various stages of development and corpora lutea could be observed in group N. In group M, the ovarian tissues displayed cystic follicles of different sizes, a reduced number of antral follicles and corpora lutea as well as an increase in the number of follicles undergoing atresia which performed nuclear pyknosis and reduced granulosa cell layer when compared with N group. Follicular development returned to normal in adolescent rats with PCOS after EA treatment, showing a significant decrease in follicular atresia and an increased number of antral follicles and corpora lutea. When compared with group M, the morphological characteristics of ovarian tissues in group SA were not significantly improved.

**Fig. 2.**
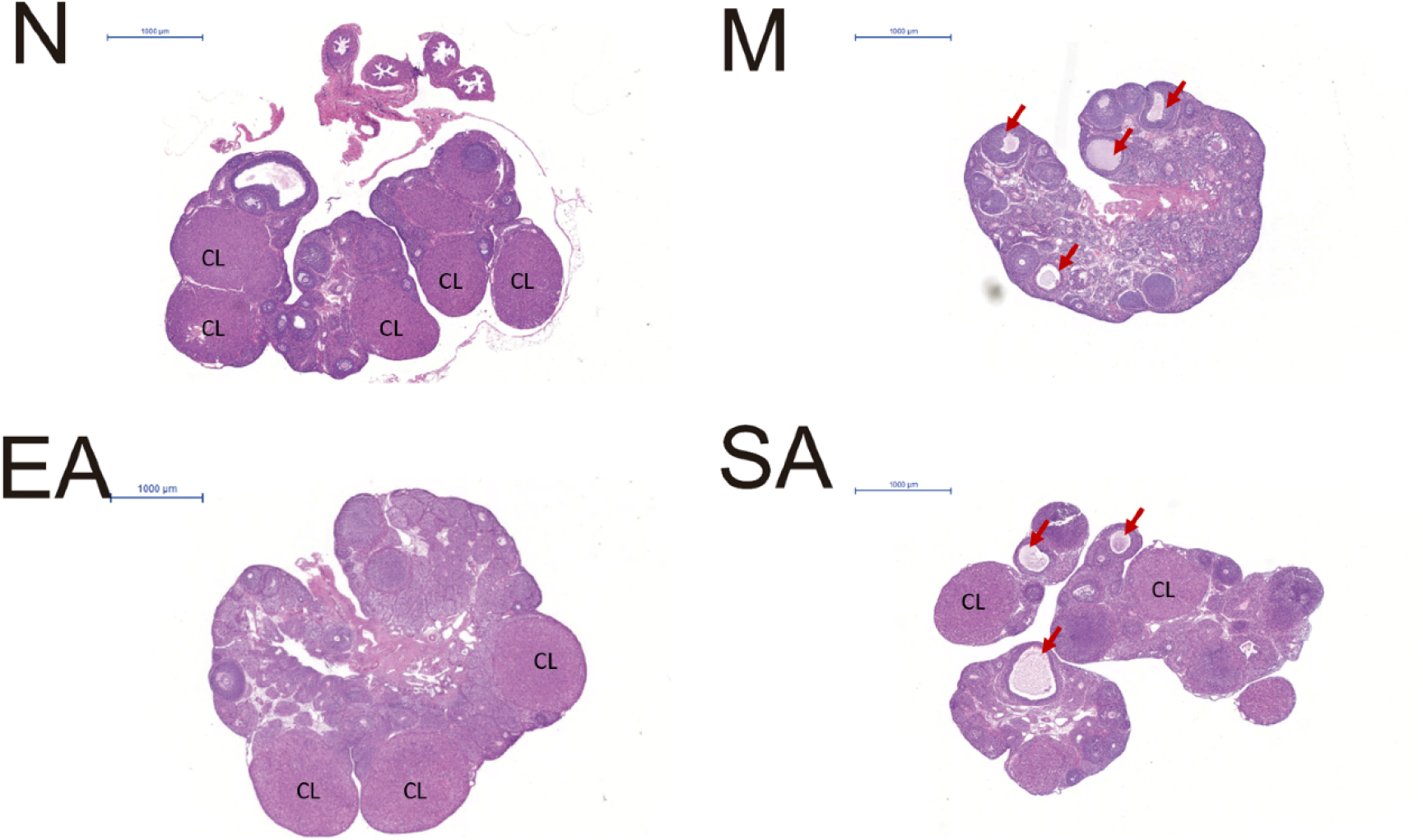
Morphological changes in rat ovarian tissues in the N, M, EA, and SA groups. N, normal group; M, model group; EA, electroacupuncture group; SA, sham acupuncture group; CL: Corpus luteum; Arrow: Cystic follicle

### Changes of body weight and serum levels of metabolic indicators

Body weight across all groups of rats showed no statistically significant (Figure 3a). Serum levels of FBG (Figure 3b) and insulin (Figure 3c) were obviously elevated in rats with PCOS when compared with normal group (both P < 0.01). EA treatment increased the level of insulin (P < 0.01) but reduced the level of FBG (P < 0.05). However, SA treatment had no significant effect. In comparison with the normal rats, the serum NPY level showed a significant increase in the model rats (P < 0.01). EA and SA both reversed this increasing level of NPY (both P < 0.01) (Figure 3d). The ghrelin level decreased in model rats (p<0.01) and could be rescued by treatment with either EA or SA (p<0.05) (Figure 3e).

**Fig. 3.**
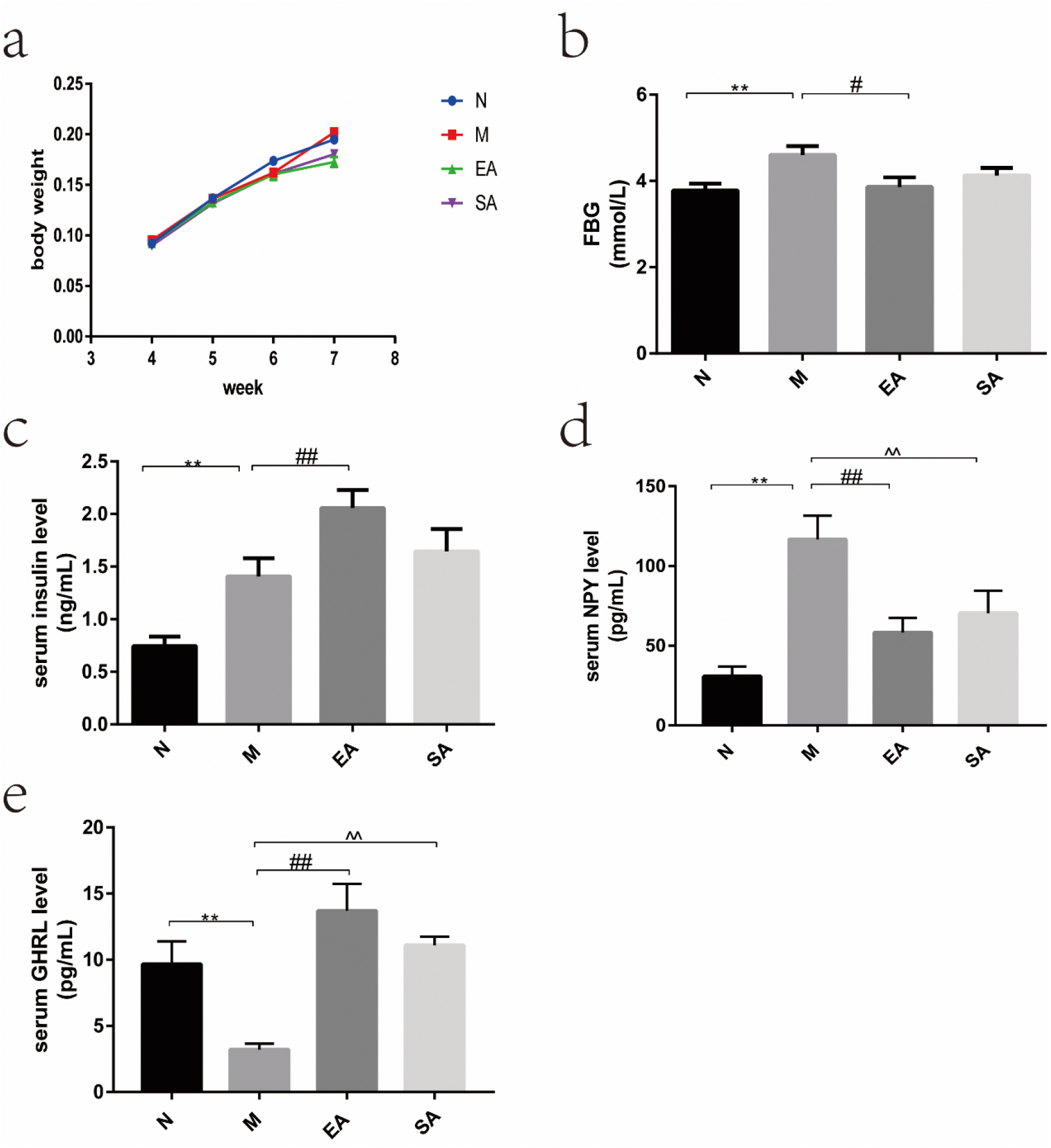
Changes of body weight and serum levels of metabolic indicators. a: body weight; b: FBG level; c: serum insulin level; d: serum NPY level; e: serum GHRL level; N, normal group; M, model group; EA, electroacupuncture group; SA, sham acupuncture group; FBG, fasting blood glucose; NPY, neuropeptide Y; GHRL, ghrelin; The values are shown as mean ± SEM. N vs M, * P < 0.05, ** P < 0.01; EA vs M, ^#^ P < 0.05, ^# #^ P < 0.01; SA vs M, ^ P < 0.05, ^ ^ P < 0.01. Eight rats per group were used in each experiment.

### Expression of GAL, GALP, GHRL and NPY in the hypothalamus

The expression of ghrelin protein (Figure 4a) and mRNA (Figure 4f) in the hypothalamus significantly decreased in the model group (both p<0.01) and EA reversed their expression (both p<0.01), while SA only reversed the mRNA expression (both p<0.01). There was no significant difference in the protein and mRNA expression of GAL and GALP between the four groups (Figure 4b-e). Unfortunately, we failed to detect the protein expression of NPY in the hypothalamus. Compared with group N, *NPY mRNA* increased significantly in the model group (p< 0.05), and both EA and SA downregulated *NPY mRNA* expression (p< 0.05) (Figure 4g).

**Fig. 4.**
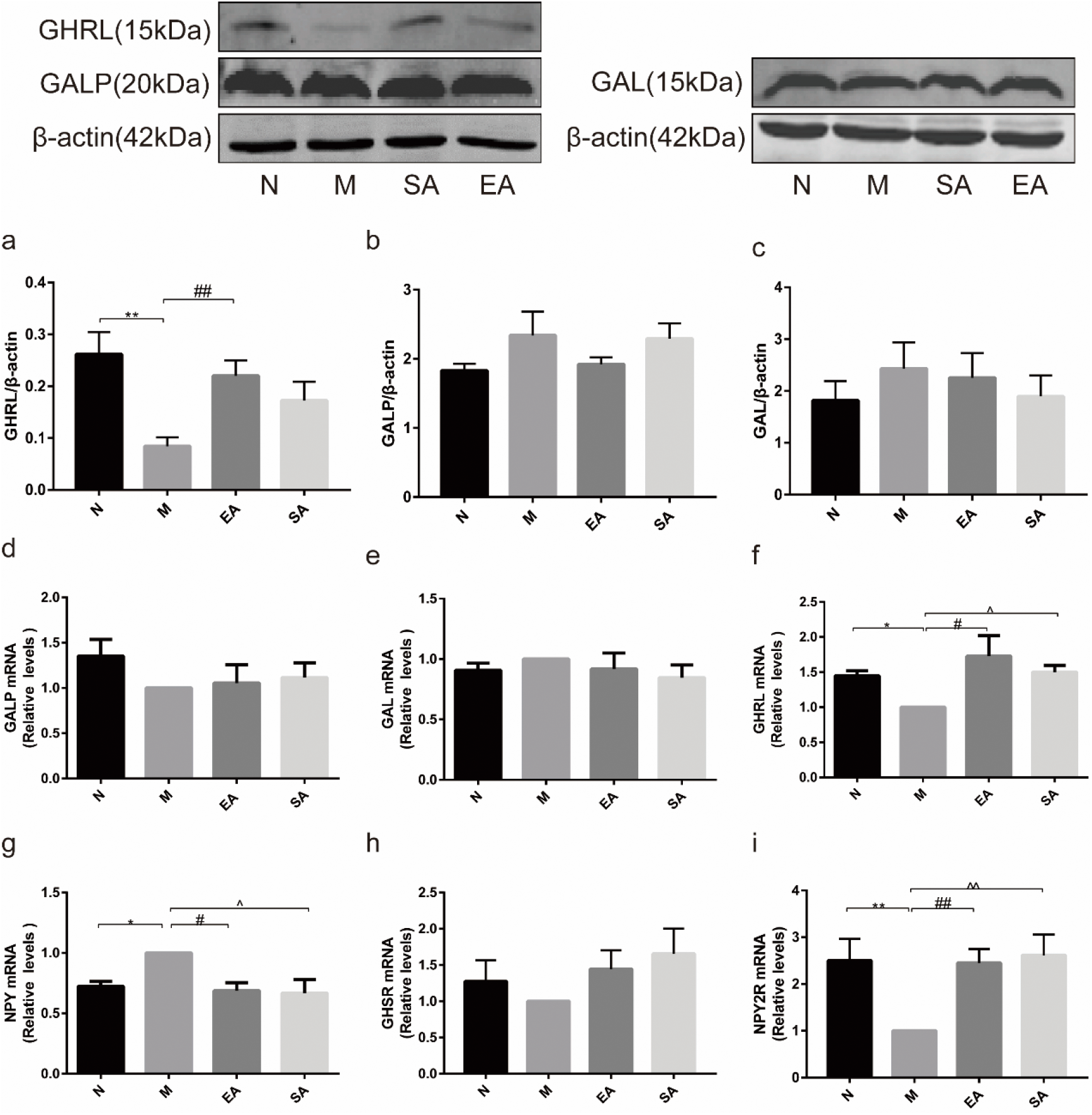
Expression of GAL, GALP, GHRL, NPY, NPY2R and GHSR in the hypothalamus. a-c: protein expression of GAL, GALP and GHRL in hypothalamus; d-i: mRNA expression of GAL, GALP, GHRL, NPY, GHSR and NPY2R in hypothalamus; N, normal group; M, model group; EA, electroacupuncture group; SA, sham acupuncture group; GAL, galanin; GALP, galanin-like peptide; NPY, neuropeptide Y; GHRL, ghrelin; GHSR, ghrelin receptor; NPY2R, neuropeptide Y2 receptor; The values are shown as mean ± SEM. N vs M, * P < 0.05, ** P < 0.01; EA vs M, ^#^ P < 0.05, ^# #^ P < 0.01; SA vs M, ^ P < 0.05, ^ ^ P < 0.01.

There were no significant differences in mRNA expression of GHSR in the four groups (Figure 4h). *NPY2R mRNA* was lower in group M than group N (P < 0.01). After treatment with EA or SA, *NPY2R mRNA* levels in rats with PCOS obviously increased (both P < 0.01) (Figure 4i).

### Expression of GHRL, NPY, NPY2R and GHSR in the arcuate nucleus (ARC)

According to the RT-PCR and WB results, we found that there were differences in the expression of NPY and GHRL in the hypothalamus. Given that the arcuate nucleus (ARC) is a key site of the reproductive regulatory system and the energy regulating system, therefore, we performed an immunohistochemistry test to determine the expression of NPY, GHRL and their receptor in the ARC. The results showed that the number of NPY-positive cells in the ARC in group M was much greater than the number in the N, EA, and SA groups (all P < 0.05), but there was no difference between these groups. Meanwhile, the number of GHRL-positive cells in the ARC in group M was much lower than the number in groups N and EA (P < 0.01, P < 0.05, respectively), but there was no difference between groups M and SA (Figure 5).There was some attenuated immunoreactivity of NPY2R in the ARC in group M compared with group N (P < 0.05), but when treated with EA or SA, the immunoreactivity of NPY2R was enhanced remarkably (P < 0.01, P < 0.05, respectively) (Figure 5).As shown by immunofluorescence, the expression of GHSR in the ARC was not significantly different in the four groups.

**Fig. 5.**
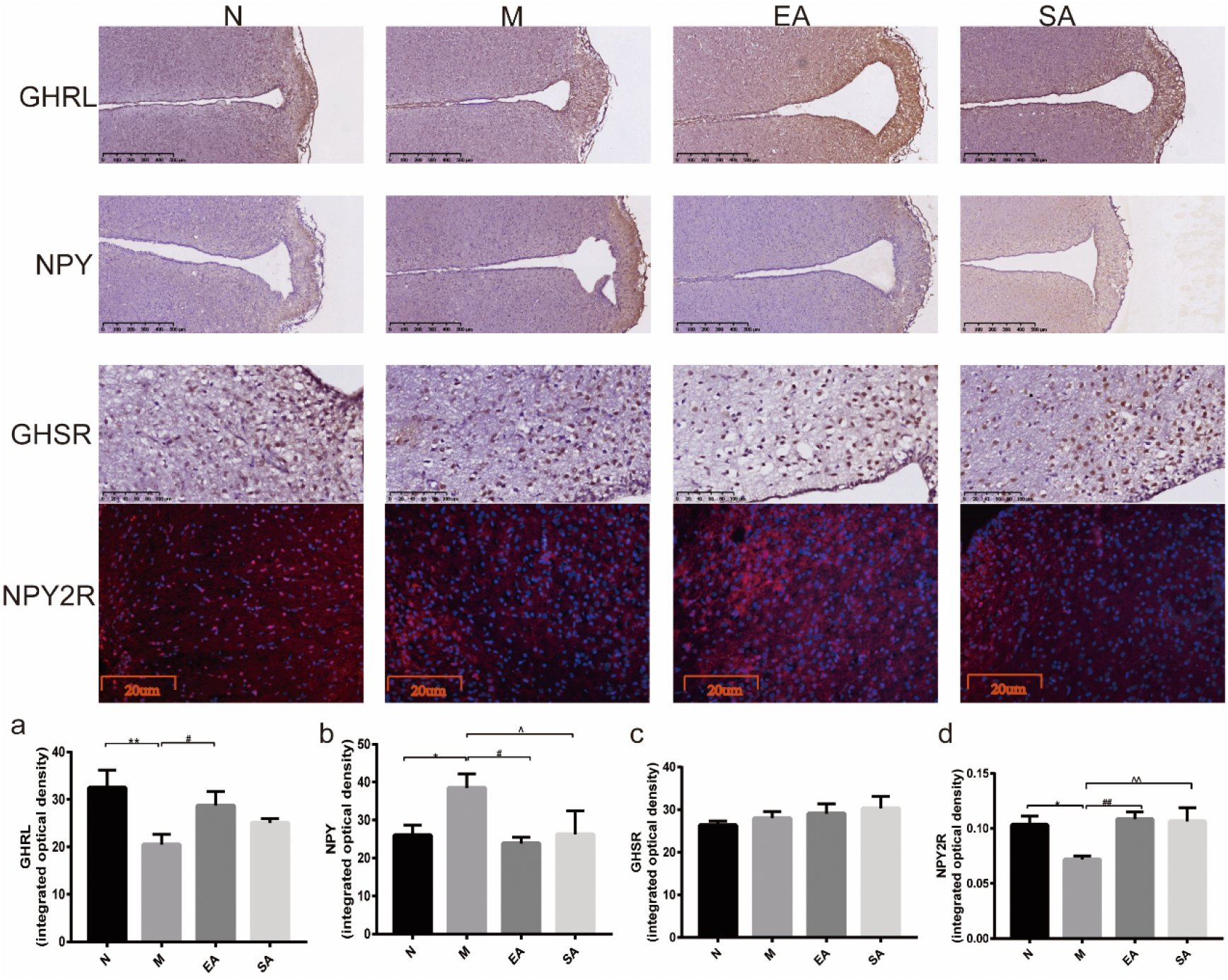
Expression of GHRL, NPY, NPY2R and GHSR in the arcuate nucleus (ARC)(a-d). N, normal group; M, model group; EA, electroacupuncture group; SA, sham acupuncture group; NPY, neuropeptide Y; GHRL, ghrelin; NPY2R, neuropeptide Y receptor; GHSR, ghrelin receptor; ARC, arcuate nucleus. The values are shown as mean ± SEM. N vs M, * P < 0.05, ** P < 0.01; EA vs M, ^#^ P < 0.05, ^# #^ P < 0.01; SA vs M, ^ P < 0.05, ^ ^ P < 0.01.

### Co-expression of NPY2R and GnRH, GHSR and kisspeptin in the ARC

As shown in Figure 6, the location of the simultaneous expression of GHSR and kisspeptin could be found in the ARC. In addition, we also observed co-expression of NPY2R and GnRH in the ARC.

**Fig. 6.**
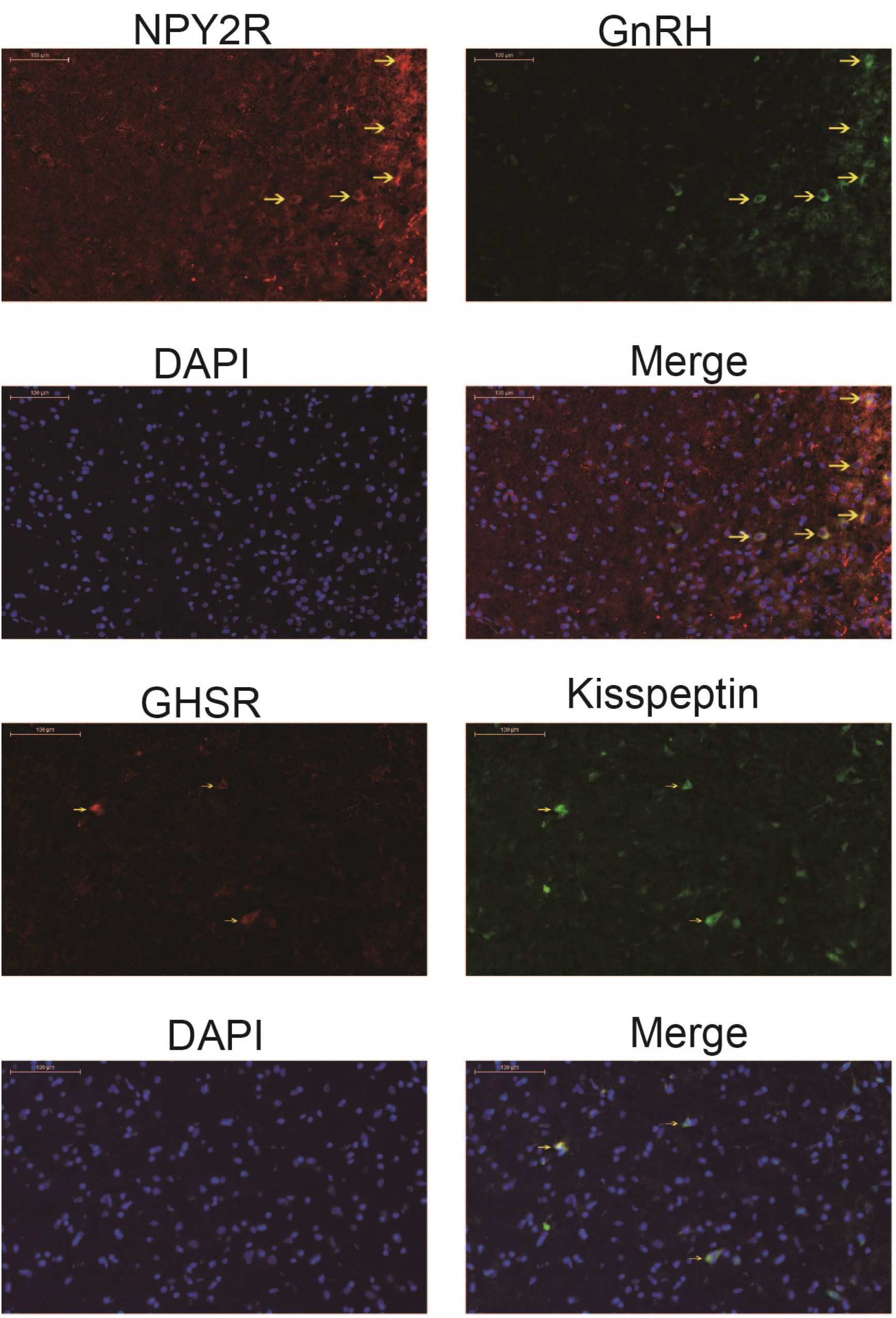
Co-expression of GHSR and Kisspeptin, as well as NPY2R and GnRH in ARC. N, normal group; M, model group; EA, electroacupuncture group; SA, sham acupuncture group; GHSR, ghrelin receptor; NPY2R, neuropeptide Y2 receptor; ARC, arcuate nucleus.

### Protein expression of kisspeptin, GALP, GAL and GHRL in the pancreas

Protein expression of kisspeptin in the pancreas increased in the model group (p<0.05). Neither EA nor SA down-regulated this increasing protein level (Figure 7a). The protein expression of GALP, GAL and GHRL did not differ between the four groups (Figure 7b-d).

**Fig. 7.**
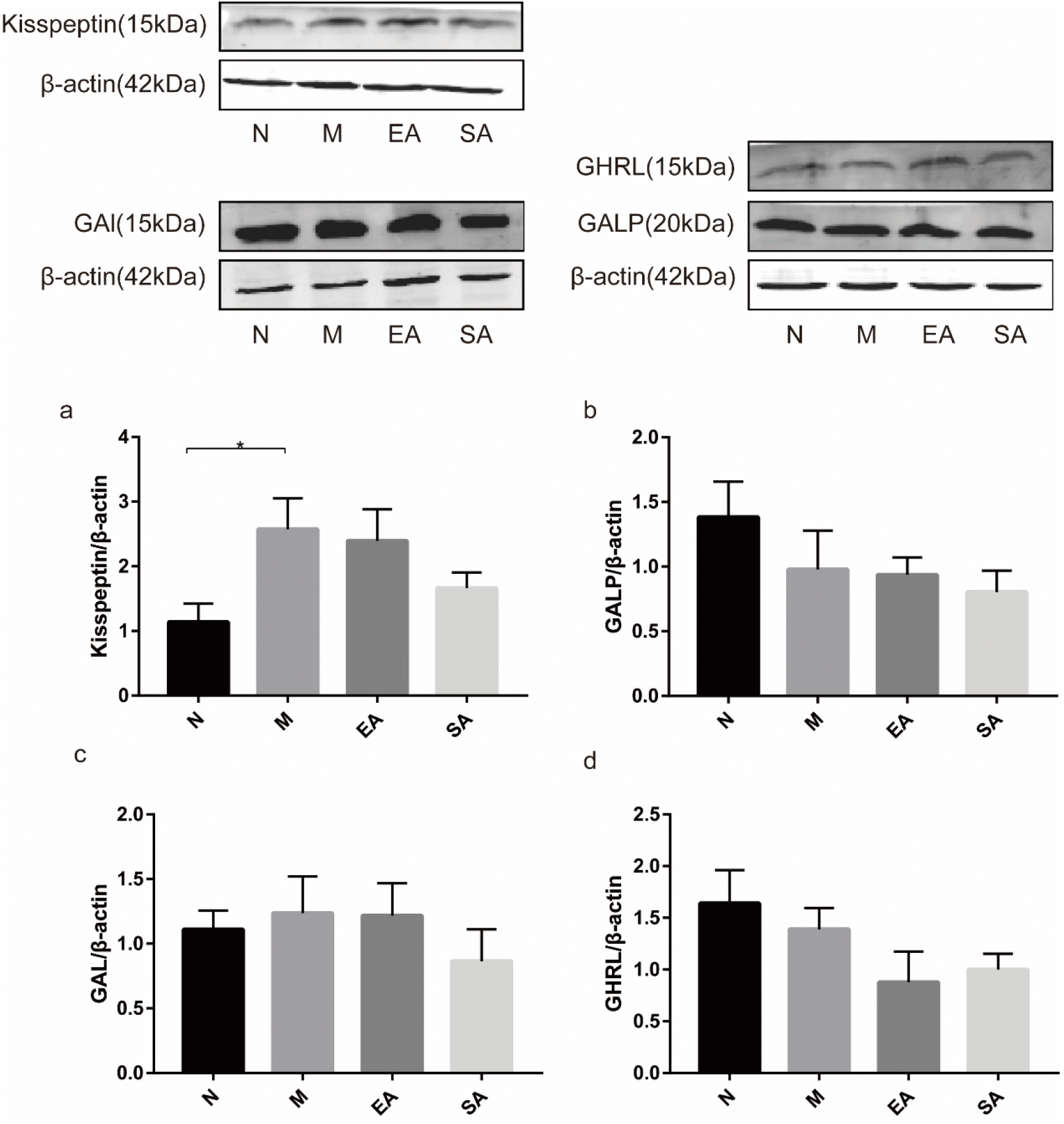
Protein expression of kisspeptin, GALP, GAL and GHRL in the pancreas (a-d). N, normal group; M, model group; EA, electroacupuncture group; SA, sham acupuncture group; GAL, galanin; GALP, galanin-like peptide; GHRL, ghrelin; The values are shown as mean ± SEM. N vs M, * P < 0.05.

## Discussion

In the present study, the pubertal rats induced with PCOS showed estrous cycle disorders and polycystic ovarian morphology. After treatment with EA, both the disordered estrous cycle and abnormal ovarian morphology were obviously improved. Meanwhile, although sham acupuncture was able to improve the disordered estrous cycle, it had no effect on ovarian morphology, suggesting that EA is significantly more effective than sham acupuncture in improving the ovarian morphology of pubertal rats with PCOS.

No significant effect on protein and mRNA expression of GAL and GALP in the hypothalamus was observed in four groups. However, protein and mRNA expression of GHRL in the hypothalamus and ARC was decreased, and NPY mRNA levels was increased in the pubertal PCOS rats, indicating that NPY and GHRL in hypothalamus are involved in the occurrence of pubertal rats with PCOS. Although the specific mechanism is unclear, we speculate that NPY and GHRL in hypothalamus may participate in development of pubertal PCOS through kisspeptin / GnRH system in view of our previous study showed that the increased expression and release of kisspeptin and GnRH in hypothalamus are the key pathogenesis of pubertal PCOS[7]. The following provides evidence for our conjecture: a, Immunofluorescence double-labeling experiment showed that GnRH neurons and kisspeptin neurons in ARC were the direct targets of NPY and GHRL in hypothalamus respectively, consistent with research of Conde et al.[24] and Dhillon et al[25]; b, Kisspeptin, as the nodal regulatory centre of reproductive function, binds into kisspeptin receptor (G-protein coupled receptor-54, GPR-54) on GnRH neurons or NPY neurons and promotes GnRH and NPY secretion from hypothalamus[26–29]; c, Previous studies have shown that NPY is necessary for the pulsatility of GnRH release[30], promoting the secretion of GnRH [10]. However, GHRL is inhibitory to GnRH/LH release[11].

The decreased GHRL mRNA levels and increased NPY mRNA levels in the hypothalamus of pubertal PCOS rats could be reversed by either EA or SA. we also observed the similar pattern in serum levels of GHRL and NPY. Furthermore, EA changed the protein expression of GHRL in the in hypothalamus and ARC of PCOS rats, and both EA and SA down-regulated NPY expression in ARC of PCOS rats. These data suggest that EA stimulation plays a beneficial role in pubertal PCOS by upregulating the expression of ghrelin and reducing the expression of NPY in hypothalamus.

It is well known that peptides play its physiological roles through a specific receptor. so we also investigated the expression of NPY and ghrelin receptors in the hypothalamus. The expression of the ghrelin receptors (formerly known as the growth hormone secretagogue receptor, GHSR) did not show any obvious differences between the four groups. Neuropeptide Y has five Y (Y1–Y5) receptor subtypes and it was found that the Y2 receptor subtype was associated with the regulation of the reproductive function by suppressing the release of GnRH and gonadotropin[31]. As expected, the expression of NPY2R in the hypothalamus of pubertal PCOS rats was clearly lower than that in normal rats in our study. Both EA and SA significantly increased the expression of NPY2R in the hypothalamus of pubertal PCOS rats.

In addition, metabolic disorders were observed in pubertal PCOS rats. In our study, pubertal PCOS rats displayed elevated FBG levels and high serum insulin levels compared with normal group. Given that kisspeptin has been shown to enhance insulin secretion from pancreas[32] and we found that the protein expression of kisspeptin obviously upregulated in pubertal PCOS rats, occurring of the increased serum insulin level in the pubertal PCOS rats may be related with the increased kisspeptin protein expression in the pancreas. EA significantly reduced FBG levels, and the underlying mechanism may be that EA promotes the production and secretion of insulin, because we found that serum insulin levels increased after EA treatment, however, EA did not change the upregulation of kisspeptin protein expression in the pancreas, indicating that kisspeptin has nothing to do with EA promoting insulin secretion. EA promoting insulin secretion and decreasing FBG levels may be relate to secretion of endogenous beta-endorphin from the adrenal gland [33, 34]. No difference in protein expression of GAL, GALP and GHRL in the pancreas were be found among four groups.

For the first time, the present research shows that EA has obvious beneficial effects on pubertal PCOS by regulating the hypothalamic expression of NPY, NPY2R, and ghrelin. However, this study seems to indicate that SA can also improve the symptoms of adolescent rats with PCOS by altering the hypothalamic expression of NPY, NPY2R, and ghrelin. One possible explanation is that SA can elicit the activity of afferent nerves in the skin, and then affect the functional connections of the brain, leading to a “limbic touch response”[35]. But SA has less impact than EA because we found that compared with EA, SA cannot change ovarian morphology and the protein expression of ghrelin in the hypothalamus.

To sum up, the expression of ghrelin decreases, which leads to increased kisspeptin levels and is accompanied by increased NPY levels in the hypothalamus, thus stimulating the overactivation of GnRH neurons and excessive GnRH secretion, ultimately resulting in the occurrence of adolescent PCOS. Stimulatory signals of EA are transmitted to the brain and play a benign regulatory role on ghrelin, NPY, and NPY2R, thereby reducing the secretion of GnRH and improving the symptoms of adolescent PCOS.

## Acknowledgments

This paper has been edited for English language, grammar, punctuation, and spelling by Enago, the editing brand of Crimson Interactive Consulting Co. Ltd. under Normal Editing.

